# Octopamine release impairs courtship conditioning in *Drosophila melanogaster*

**DOI:** 10.1101/034827

**Authors:** Julia Chartove, Eddie Zhang, Cici Zhang

## Abstract

Octopamine is known to have an appetitive role in odor conditioning paradigms in Drosophila melanogaster. We wanted to test whether octopamine could also act as an appetitive stimulus in courtship conditioning, a paradigm in which training with an unreceptive female (such as a decapitated virgin) causes a subsequent decrease in courtship behavior in male Drosophila. To control octopamine release, we used the Tdc2-Gal4 and UAS-dTRPa1 genes in conjunction to depolarize octopaminergic neurons at 27 C in experimental flies. We hypothesized that inducing octopamine release during courtship training would decrease the aversive impact of training and cause less subsequent suppression of courtship behavior. Our findings confirmed this hypothesis: Tdc2-Gal4/UAS-dTRPa1 flies trained at 27 degrees showed significantly more courtship behavior than controls during testing, and in fact showed no significant effect of courtship training. This confirms that octopamine release counteracts the aversive stimulus of failure to copulate, indicating that octopamine may have an appetitive role in courtship.

## Introduction

Octopamine is a monoamine neurotransmitter that plays an important role in many invertebrate systems. It is structurally very similar to norepinephrine with the only difference being that norepinephrine contains an additional hydroxyl group on the central benzene ring. Therefore, octopamine has been likened to an invertebrate analogue of norepinephrine (Blenau and Baumann, 2001). In *Drosophila melanogaster*, octopamine plays a role in many functions such as locomotion, courtship, and fertility (Chen *et al*, 2012). It is necessary for male-male aggression (Hoyer *et al*, 2008) and modulates the decision between courtship and aggression towards other males (Certel *et al*, 2010). Octopamine is also involved in memory, specifically appetitive memory formation. Octopamine is required for the formation of olfactory conditioning associated with an appetitive stimulus (sugar), but not required for olfactory conditioning associated with an aversive stimulus (shock) (Schwaerzel *et al*, 2003). In addition, stimulation of octopaminergic neurons in association with a scent is sufficient to produce appetitive memory (Schroll *et al*, 2006). This indicates that octopamine acts as a reward hormone that encodes positive valence of a stimulus.

Octopamine and dopamine encode opposite valence in odor conditioning paradigms in *Drosophila*, with octopamine encoding a positive stimulus and dopamine encoding a negative stimulus. These two neurotransmitters act within the same anatomical regions to produce both aversive and appetitive memory (Schwaerzel *et al*, 2003). Similarly opposed effects of octopamine and dopamine have been observed in many invertebrate species. In the honeybee, octopamine enhances appetitive odor memory formation, while dopamine impairs its retrieval (Scheiner *et al*, 2006). In the cricket, octopamine is necessary for appetitive but not aversive taste conditioning, while the opposite is true for dopamine (Unoki *et al*, 2005). The same is true of the crab with regards to visual learning; furthermore, injection of dopamine impairs the formation of appetitive memory, while injection of octopamine impairs aversive memory formation (Klappenbach *et al*. 2012, Kaczer and Maldonado 2009). This body of research indicates that the opposite valences of octopamine and dopamine are robust across species and training paradigms, and that the octopamine and dopamine circuits are likely capable of interacting in opposition to each other to determine the overall valence of a stimulus. Therefore, we can expect to observe similar interactions between these two neurotransmitter systems in other learning paradigms in *Drosophila*, such as (in our experiment) courtship conditioning.

The courtship conditioning behavioral assay is an established paradigm for assessing aversive learning in *Drosophila*. Courtship suppression is induced by pairing a male fly with a sexually unreceptive female for an extended period. After this training, the male will show reduced courtship behavior towards new females (Griffith 2009). Keleman *et al* (2012) demonstrated that dopamine is necessary for courtship suppression, and furthermore that inducing dopaminergic neural activity in naive males can induce courtship suppression in the absence of training. As dopamine can mimic the aversive effects of courtship conditioning, octopamine could potentially have an appetitive influence on courtship behavior. If this is the case, then octopamine release during courtship training should counteract the negative valence encoded by the dopamine circuit during memory formation, as seen in Kaczer and Maldonado (2009). We hypothesize that due to octopamine’s appetitive valence, elevated octopamine activity during courtship training could impair aversive courtship memory formation.

To induce octopaminergic activation in this experiment, we utilized the Gal4-UAS system to cause expression of the *UAS-dTrpA1* effector, which encodes for a heat-sensitive ion channel that opens at 25 C (Pulver *et al* 2009), in octopaminergic neuron populations indicated by the *Tdc2-Gal4* driver (Certel *et al* 2010). This gene system causes octopaminergic neurons in flies expressing both Tdc2-Gal4 and UAS-dTRPa1 to depolarize at high temperatures. However, Tdc2-Gal4 or UAS-dTRPa1 alone do not have any phenotypical effect; therefore, flies only expressing one transgene or the other may be used as genetic controls. Since the dTRPa1 channel only opens at temperatures above 25 C, Tdc2-Gal4/UAS-dTRPa1 flies raised and trained at lower temperatures are phenotypically identical to wild type flies and may be used as temperature controls. Though a large proportion of prior research has used mated females in inducing courtship suppression, this study used decapitated virgin females in both training and testing. This was done to narrow down the causality of the courtship suppression effect to the failure to copulate during training, excluding the aversive stimulus of a mated female’s pheromone profile (Ejima 2005). Our hypothesis predicts that aversive courtship memory formation will be impaired in experimental populations of flies expressing both the *UAS-* and *Gal4-*transgenes trained at 27 C with decapitated virgins, and therefore that these flies will exhibit less courtship suppression than genetic or temperature controls.

## Materials and Methods

### Drosophila strains and genetic crosses

All flies were fed a cornmeal, sugar, and agar food mixture, the preparation of which is detailed in the Siwicki Lab’s Food Protocol. *Tdc2-Gal4, UAS-dTrpA1*, and Canton-S (wild type) fly stocks were raised on a 12 hourr light, 12 hour dark cycle at 25 C at constant humidity. All flies were collected and sorted by sex into separate vials of 10 flies each within 4–6 hours of lights-on to ensure virginity. Any single-sex vial that contained larvae was discarded, as this indicated that the virginity of the flies within the vial had been compromised. Parent flies were transferred into new vials every 7 days to expand stocks. After three weeks of stock expansion, ~10 male and ~10 female flies each were transferred to new breeding vials and crossed to produce control (containing either the *Uas-* or *Gal4-*transgene) and experimental (expressing both *Uas-* and *Gal4-*transgenes) genotypes. *UAS-dTrpA1* flies were crossed with Tdc2-Gal4 flies, creating F1 offspring with the *Tdc2-Gal4/ UAS-dTrpA1* experimental genotype. *UAS-dTrpA1* and *Tdc2-Gal4* flies were crossed with wild-type flies to create the control genotypes *UAS-dTrpA1/+* and *Tdc2-Gal4/+*, in which there were either no *Gal4* transgene to drive expression or no *UAS* transgene to encode the temperature-sensitive *dTrpA1* ion channel. Experimental and genetic control flies were all raised at 19 C, a temperature at which the *dTrpA1* ion channel is inactive, to control for any developmental differences potentially caused by temperature or elevated octopamine activity. Wild-type Canton-S flies used for crosses and as female trainers and testers were reared in a separate incubator at 25 C.

### Short-term memory training procedure

All flies (male and female) were aged 4–7 days before experimental use. Socially naive male flies were trained in either a 19 C or 27 C incubator for one hour either with a decapitated virgin (courtship training) or in isolation (sham training). (See table 1 for a full list of experimental and control groups.) These temperatures were chosen because at 27 C, the dTrpA1 ion channel is active, elevating octopamine levels in Tdc2-Gal4/UAS-dTRPa1 flies; at 19 C, it is inactive, keeping octopamine levels at baseline. After training for one hour, trained males were removed from incubators and allowed to recover from temperature variations for ten minutes. Any trainer females were then discarded and all males were transferred to chambers containing new decapitated virgins for testing, all of which was conducted at 19 C. Behavior was recorded for ten minutes after the pairing of each male with decapitated female.

**Table 1.** Experimental and control groups by genetic manipulation, training temperature, and training condition. dTRPa1 is only active in the experimental group and the sham control; the other flies exhibit wild-type phenotypes.

| Group | Genotype | Temperature during training | Trainer fly | n |
| --- | --- | --- | --- | --- |
| Experimental | Tdc2-Gal4/UAS-dTRPa1 | 27 C | Decapitated virgin | 20 |
| Sham control | Tdc2-Gal4/UAS-dTRPa1 | 27 C | No fly (sham) | 20 |
| Temperature control | Tdc2-Gal4/UAS-dTRPa1 | 19 C | Decapitated virgin | 15 |
| Sham temperature control | Tdc2-Gal4/UAS-dTRPa1 | 19 C | No fly (sham) | 13 |
| Genetic control | Tdc2-Gal4/+ or UAS-dTRPa1/+ | 27 C | Decapitated virgin | 19 |
| Sham genetic control | Tdc2-Gal4/+ or UAS-dTRPa1/+ | 27 C | No fly (sham) | 19 |
| Genetic temperature control | Tdc2-Gal4/+ or UAS-dTRPa1/+ | 19 C | Decapitated virgin | 9 |
| Sham genetic temperature control | Tdc2-Gal4/+ or UAS-dTRPa1/+ | 19 C | No fly (sham) | 11 |

#### Data Analysis and Scoring

Scorers who were blind to genotype and training conditions scored recorded videos for number of bouts of courtship behavior and latency of courtship initiation after pairing. The total number of observations per fly varied only slightly across scorers, indicating consistency of scoring technique. Analysis was performed by examining each fly for a two-second interval every ten seconds. Behaviors considered to be courtship were limited to tapping, winging, licking, and attempted copulation. Typical courtship behavior such as following was not included due to the immobility of decapitated females, and inactive courtship behaviors like orientation of the male toward the female or idling by the female were also not included. Latency to initiate courting and number of courting observations were used to calculate courtship index (CI, proportion of time spent courting weighted by courtship initiation latency):

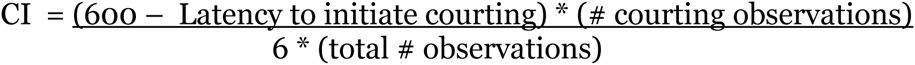

Data analysis was performed by conducting t-tests between groups. *UAS-dTrpA1* and *Tdc2-Gal4* genetic controls did not exhibit a significant difference (p>0.05) in CI at either 19C or 27C and were collapsed into a single control genotype group for greater statistical power during analysis.

## Results

As hypothesised, courtship conditioning was impaired in Tdc2-Gal4/UAS-dTRPa1 experimental genotype flies trained with decapitated virgins at 27 C. These flies showed a significantly higher courtship index during testing than genetic controls trained at 27 C (t(n = 39), d.f. = 37, p = 0.0382), indicating that the courtship memory was weaker in experimental flies. Additionally, though courtship conditioning had a significant effect in both genetic and temperature controls, such an effect was not seen in the experimental group. Both genetic controls trained at 27 degrees (t(n = 38), d.f. = 33, p = 0.0009) and Tdc2-Gal4/UAS-dTRPa1 flies trained at 19 degrees (t(n = 28), d.f. = 20, p = 0.0192) showed a significantly lower courtship index than sham trained groups. However, no significant effect was observed in the experimental group (t(n = 40), d.f. = 36, p= 0.1512), which suggests that little to no courtship memory was induced in the experimental flies.

Results from control groups suggest that the observed impairment in courtship conditioning was due solely to the temperature-dependent effects of Tdc2-Gal4/UAS-dTRPa1. No effects of temperature during training were found on genetic controls in either the trained condition (t(n = 28), d.f. = 14, p = 0.7572) or the sham condition (t(n = 30), d.f. = 22, p = 0.6541). This confirms that temperature does not have an effect on courtship memory formation in flies expressing a normal phenotype. Additionally, Tdc2-Gal4/UAS-dTRPa1 flies trained with decapitated females at 19 C do not have a significant difference in mean courtship index from genetic controls trained under the same conditions. (t(n = 24), d.f. = 13, p = 0.9466). Furthermore, the difference in courtship index between Tdc2-Gal4/UAS-dTRPa1 flies trained with decapitated virgins at 27 C and at 19 C is very nearly significant (t(n = 35), d.f. = 33, p = 0.0723). Taken together, these results indicate that the dTRPa1 channel was only active in Tdc2-Gal4/UAS-dTRPa1 flies at 27 C, and therefore that the genetic and temperature manipulations only affected the experimental group. Because all male flies used in this experiment were raised at 19 C, we can safely conclude that the dTRPa1 channel was not active in the Tdc2-Gal4/UAS-dTRPa1 flies at any point before testing. Therefore, it is reasonable to assume that there were no developmental differences between experimental flies and genetic controls.

## Discussion

In this study, Tdc2-Gal4/UAS-dTRPa1 experimental flies showed significantly more courtship behavior than both temperature controls and genetic controls when trained with decapitated females at 27 degrees, a temperature at which the dTRPa1 ion channel induces elevated firing in octopaminergic neurons. (Figure 1) These results confirmed our hypothesis that octopamine release disrupts aversive courtship conditioning, possibly via acting as an appetitive stimulus. Moreover, the octopamine effects we observed are not due to inherent effects of temperature or to developmental differences between Tdc2-Gal4/UAS-dTRPa1 flies and genetic controls. We saw no effects of temperature in genetic controls (Figure 3), nor was there a difference between controls and Tdc2-Gal4/UAS-dTRPa1 flies trained at 19 degrees (Figure 4). This indicates that the impaired conditioning effects we observed were solely due to octopamine release during training. It is interesting to note that not only does octopamine reduce the effects of courtship conditioning, it almost completely counteracts it: the effect of courtship conditioning on Tdc2-Gal4/UAS-dTRPa1 flies trained at 27 degrees is insignificant (Figure 2).

**Figure 1.**
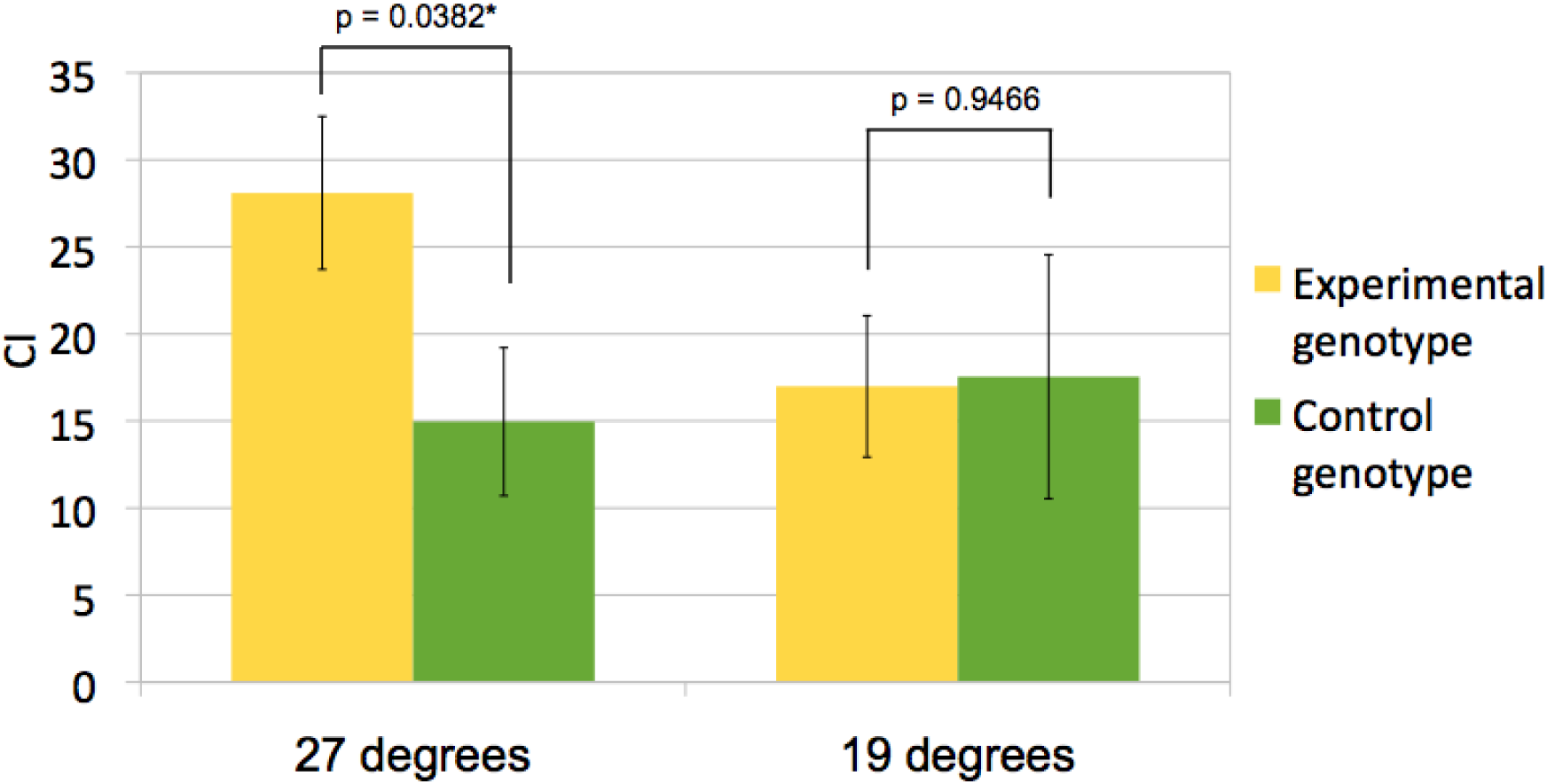
Courtship index after training with decapitated females in Tdc2-Gal4/UAS-dTRPa1 flies versus genetic controls. Significantly more courtship behavior was observed in Tdc2-Gal4/UAS-dTRPa1 flies at 27 degrees, indicating that aversive courtship conditioning was impaired at that temperature due to octopaminergic effects of dTRPa1 during training.

**Figure 2.**
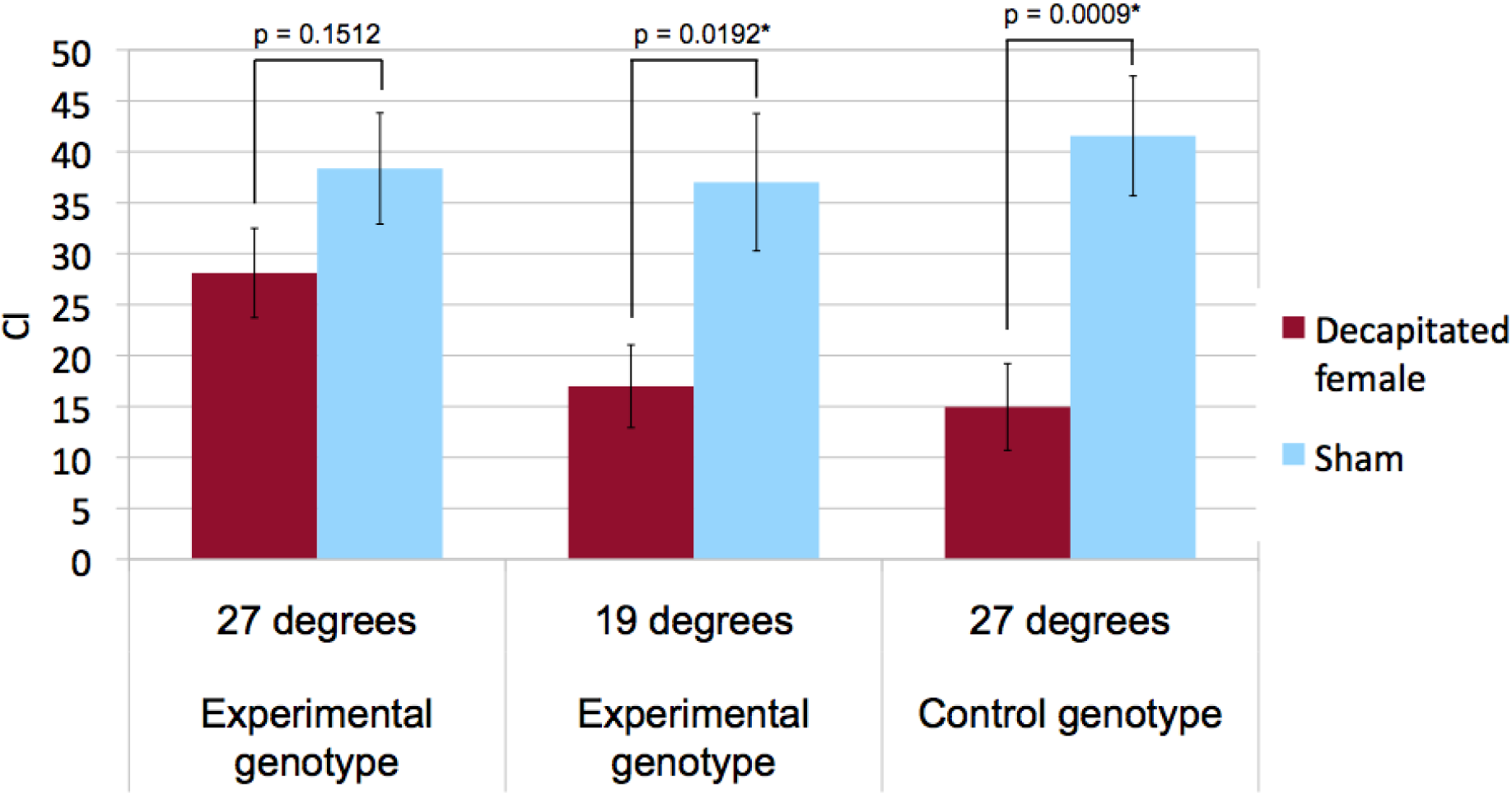
Effects of training with decapitated females in experimental and control groups. Genetic and temperature control groups show significant decreases in courtship behavior due to training, whereas the experimental group does not. This indicates that octopamine release almost completely counteracts courtship suppression.

**Figure 3.**
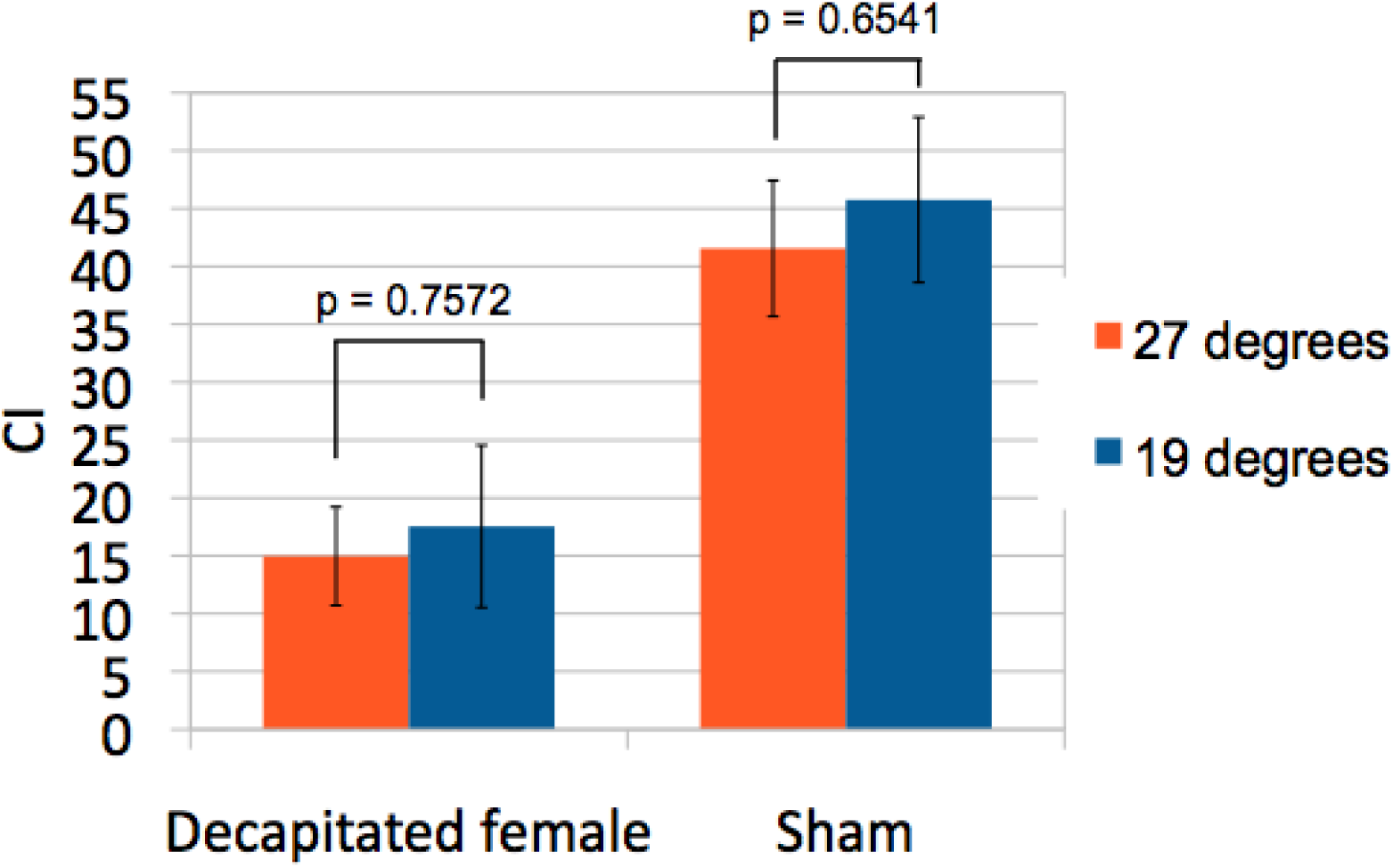
Courtship index at different temperatures in genetic controls. Both temperature groups showed a similarly low courtship index after training, indicating that there are no effects of temperature on courtship conditioning in normal flies.

**Figure 4.**
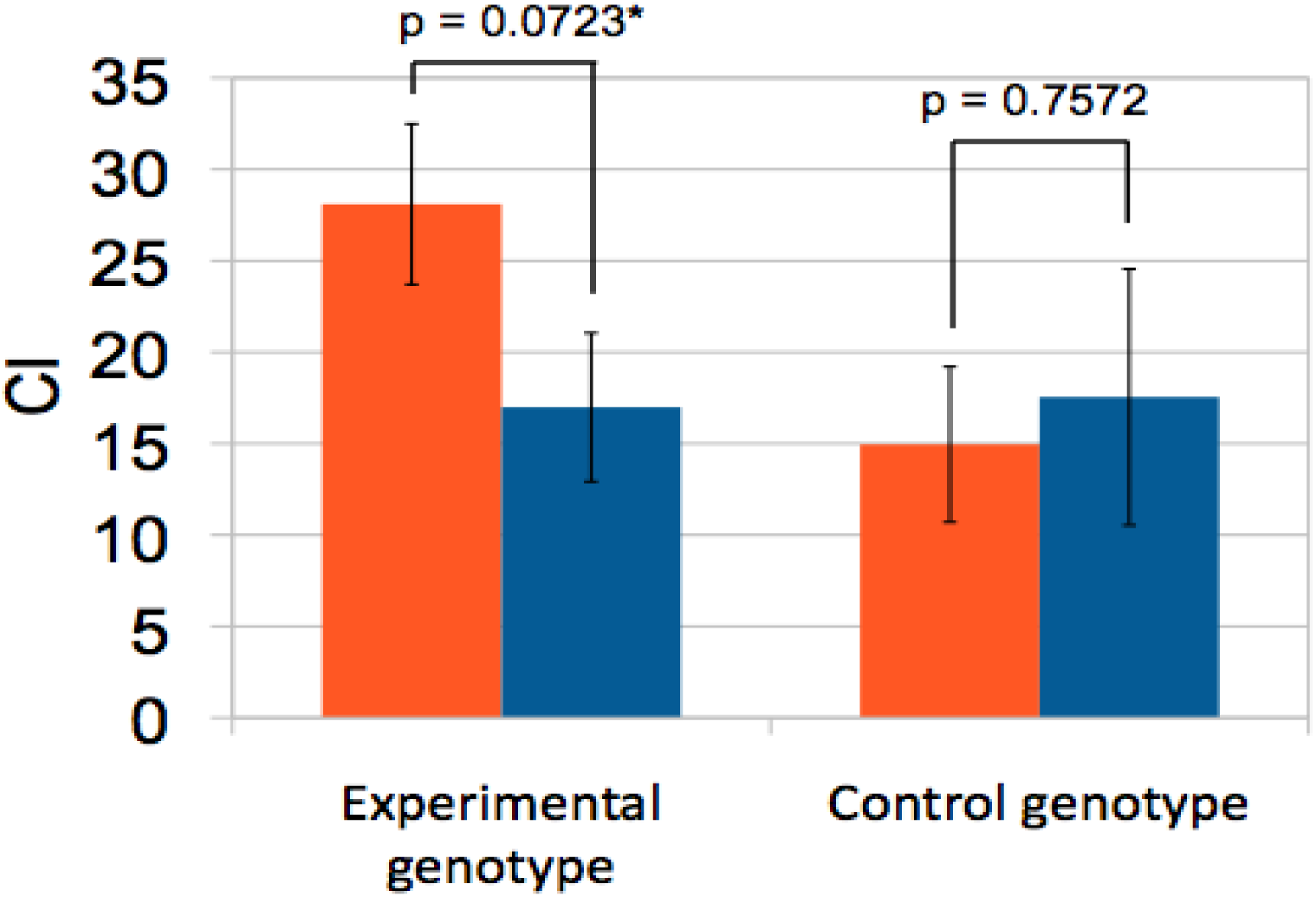
Experimental flies and genetic controls trained with decapitated females. Temperature effects approach significance for the experimental group, suggesting that dTRPa1 was inactive at 19 degrees. As experimental flies were raised at 19 degrees, we can assume they experienced no effects of dTRPa1 during development.

Previous studies in *Drosophila* have established octopamine as an appetitive stimulus in olfactory conditioning paradigms (Schroll et al, 2006; Schwaerzel et al, 2003). However, ours is the first to discover that octopamine release has effects on courtship conditioning. Although our results demonstrated that octopamine disrupts aversive courtship conditioning, the mechanism underlying this phenomenon remains unknown. It is conceivable that octopamine may impact courtship conditioning by acting as an appetitive stimulus. Given the opposing effects of octopamine and dopamine found in different animal models and training paradigms (Kaczer and Maldonado, 2009; Schwaerzel et al, 2003; Unoki et al, 2005), and because dopamine release is believed to be necessary for aversive conditioning (Klappenbach et al, 2012; Schwaerzel et al, 2003; Unoki et al, 2005), we speculate that octopamine may impair courtship conditioning by interacting with the dopamine circuit. One potential way to study such interaction is to apply high-performance liquid chromatography (HPLC) to measure and compare the levels of neurotransmitters in extracts of fly brains (McClung and Hirsh, 1999). If it were the case that octopaminergic activity prevents dopamine release, we could observe significant lower dopamine levels in Tdc2-Gal4/UAS-dTRPa1 flies trained at 27 C than in controls. Other, more complex interactions between these two circuits could be discovered as well.

It is also not known precisely what role octopamine plays in courtship on the whole. Previous studies have demonstrated that flies lacking both octopamine and dopamine do not display courtship behavior at all, whereas the presence of either neurotransmitter is sufficient to allow courtship behavior to occur (Chen et al, 2012). These findings suggest that octopamine probably plays a role in courtship behavior in wild-type *Drosophila*, and that it likely interacts with the dopamine circuit while doing so. Since we observed that octopamine release can counteract the aversive stimulus of failure to copulate, it may be the case that octopamine is released during successful copulation in flies, and therefore encodes an appetitive stimulus. However, we cannot conclude from our research that octopamine necessarily has an appetitive role in courtship; it may impair courtship memory through some other mechanism that disrupts memory function.

In addition, previous research suggests that octopamine’s role in courtship conditioning may go beyond simply impairing memory formation at above-baseline levels. K.M.C. O’ Dell (1994) found that flies carrying the *inactive* allele, which only have 15% as much octopamine as is found in wild-type flies, show impaired courtship conditioning. In addition, Zhou et al (2012) showed that deactivating octopaminergic neurons or tyramine β hydroxylase, the enzyme responsible for synthesizing octopamine, caused similar impairment of courtship conditioning. Considering that evidence together with our study, in which high octopamine levels impaired courtship conditioning, one could imagine a dose-dependent effect of octopamine on the formation of courtship memory: either too little or too much octopamine would have a negative impact. This suggests a regulatory role for octopamine in courtship learning rather than a straightforward positive valence.

Despite our significant and robust results, our experimental procedure still had potential sources of error which could be improved in future experiments. These include controlling for humidity and for each subject’s resting time between training and testing, neither of which were recorded during our experiment. However, we are confident that more important factors, such as consistency of video scoring and temperature conditions, were adequately controlled for. Although each fly was scored by one of three individuals, scorers trained and practiced together following the standardized Siwicki lab protocol. The slight temperature fluctuations during training due to occasional opening of the incubator should not have made any major difference on the amount of octopamine release in experimental genotype flies: the door was only open for seconds, which should not have had a significant impact on the one-hour training sessions.

In addition to the dopamine assay suggested above, several potential follow-up experiments could be conducted based on our research. All testing in this experiment was conducted at 19 C; if the experiment were repeated with tests conducted at 27 C, we could explore whether octopamine release impairs aversive memory retrieval as well as formation. It would also be interesting to see the effects of training and testing using mated females as opposed to decapitated virgins, despite the fact that previous studies have found similar courtship suppression effects for both (Keleman *et al* 2012). Theoretically, the aversive stimulus in courtship conditioning is the failure to copulate, so the type of trainer would not matter. However, the aversive pheromonal profile of mated females might induce stronger courtship memory due to the presence of additional olfactory cues (Siwicki et al, 2005). Therefore, experiments using mated females as well as decapitated mated females in order to combine the effects of immobility and aversive pheromones would inform our current findings. We could also examine whether we would observe similar results using trainers and testers of different types in order to determine whether the effects we observed are true of more generalized memory. Another potential question to research is whether octopamine is released during copulation, as suggested above. This could be addressed using HPLC to assess octopamine levels in male flies immediately after copulation.

## Acknowledgements

We would like to thank Kathleen Siwicki for her guidance in developing our experimental design, provision of lab equipment, and instruction in experimental protocols. We would also like to thank Nick Kaplinsky for allowing us to use his incubator.

